# Extracellular Vesicles from *Pneumocystis carinii*-Infected Rats Impair Fungal Viability but are Dispensable for Macrophage Functions

**DOI:** 10.1101/2023.09.19.558454

**Authors:** Steven G. Sayson, Alan Ashbaugh, Melanie T. Cushion

## Abstract

*Pneumocystis* spp. are host obligate fungal pathogens that can cause severe pneumonia in mammals and rely heavily on their host for essential nutrients. The lack of a sustainable *in vitro* culture system poses challenges in understanding their metabolism and the acquisition of essential nutrients from host lungs remains unexplored.

Transmission electron micrographs show Extracellular Vesicles (EVs) are found near *Pneumocystis* spp. within the lung. We hypothesized that EVs transport essential nutrients to the fungi during infection. To investigate this, EVs from *P. carinii* and *P. murina* infected rodents were biochemically and functionally characterized. These EVs contained host proteins involved in cellular, metabolic, and immune processes as well as proteins with homologs found in other fungal EV proteomes, indicating *Pneumocystis* may release EVs. Notably, EV uptake by *P. carinii* indicated their potential involvement in nutrient acquisition and indicate a possibility for using engineered EVs for efficient therapeutic delivery. However, EVs added to *P. carinii in vitro*, did not show increased growth or viability, implying that additional nutrients or factors are necessary to support their metabolic requirements. Exposure of macrophages to EVs increased proinflammatory cytokine levels, but did not affect macrophages’ ability to kill or phagocytose *P. carinii*. These findings provide vital insights into *P. carinii* and host EV interactions, yet the mechanisms underlying *P. carinii*’s survival in the lung remain uncertain. These studies are the first to isolate, characterize, and functionally assess EVs from *Pneumocystis*-infected rodents, promising to enhance our understanding of host-pathogen dynamics and therapeutic potential.

## Introduction

*Pneumocystis* spp. are host obligate fungal pathogens that can cause a lethal pneumonia in humans and other mammals. Most species only infect a single mammalian species and the species infecting humans is *Pneumocystis jirovecii* with the resultant pneumonia called *<u>Pneumocystis</u> jirovecii* <u>p</u>neumonia (PJP). In rats, a model used in the present study, the pneumonia is termed *<u>Pneumocystis carinii</u>* <u>p</u>neumonia (PCP). These fungi are extracellular, stenoxenous parasites that reside in the lung alveoli of mammals. Within the alveolar lumen, *Pneumocystis* spp. are thought to produce a biofilm-like system (1) with individual organisms forming a tight interdigitation with alveolar epithelial type I (ATI) cells, serving as an anchor for the pathogen clusters. Besides the fungal organisms, components of the *Pneumocystis* spp.-infected alveoli revealed by Transmission Electron Micrographs (TEMs) include lamellar bodies, tubular myelin, and notably, double-membraned vesicles (2, 3). These vesicles that have been thought to be filopodia serving for attachment, but there is disagreement as to their form and function, and no studies have been conducted to understand their purpose.

Characteristic of host-obligate pathogens, *Pneumocystis* spp. have a highly compact genome with considerable loss of many biological pathways, making them highly dependent on the host for nutrients (4, 5). These fungi have limited capacity for synthesizing amino acids, with only 2 enzymes present in the genome that are required for the synthesis of amino acids, which is insuffiecient for *de* novo synthesis of any of the 20 amino acids (4, 5). In contrast, *Schizosaccharomyces pombe* (a phylogenetically close relative) contains all 54 enzymes required for amino acid synthesis. Additionally, *Pneumocystis spp.* do not contain biochemically detectable ergosterol, the major sterol contained within the cell membrane of most fungi. Cholesterol is the most prominent bulk sterol in *Pneumocystis* spp. and is considered to be transferred from the host (6–8). However, the mechanism of amino acid and cholesterol acquisition, as well as other nutrients, by *Pneumocystis* spp. remain unclear.

It is our contention that the long-observed vesicles accompanying *Pneumocystis* spp. infections are Extracellular Vesicles (EVs) that serve in part, to shuttle essential nutrients that these fungi can no longer synthesize.

Fungal EVs have been described in terms of secretion, biological contents, and intercellular communication. Several fungi, such as *Cryptococcus neoformans* (9), *Histoplasma capsulatam* (10), *Paracoccidioides brasiliensis* (11), *Malassezia sympodialis* (12), *Saccharomyces cerevisiae* (13), *and Aspergilus fumigatus* (14), have shown secretion of EVs. Like those in mammals, these vesicles contain a diverse composition of proteins, lipids, nucleic acids, and polysaccharides (15). *Candida albicans* secretes EVs that participate in community communication and are required for the proper formation of biofilms (16). Although the uptake of fungal EVs by host cells has been documented (17), there is limited evidence demonstrating the uptake of host EVs by fungal organisms.

In the mammalian lung, alveolar epithelial type I (ATI) cells are responsible for gas exchange with the capillaries and are the first line of defense against inhaled stimuli. In response to stimuli, ATI cells release extracellular vesicles (EVs) which modulate the lung environment (18, 19). These mammalian EVs have been shown to contain DNA, RNA, and protein components that are involved in intercellular communication (20). Additionally, cholesterol and free amino acids have been found to be enriched in exosome and microvesicles from most tissues and cell types, including lungs, epithelial cells, and macrophages (21–23).

In the present work, nano-scale liquid chromatographic tandem mass spectrometry (nLC-MS/MS) analysis was used to characterize the proteome of bronchial alveolar lavage fluid (BALF)-derived EVs from *P. carinii*-infected rats and *P. murina*-infected mice. The *Pneumocystis-*specific Major surface glycoprotein (Msg), as well as other *Pneumocystis* proteins, were detected, indicating the potential source(s) of these EVs. These proteomic profiles supports the hypothesis that *P. carinii* and *P. murina* may themselves release a population of EVs while uptake studies revealed that *P. carinii* can uptake host-derived EVs.

## Results

### P. carinii-infected rat BALF contain EVs

The presence of EVs within *P. carinii*-infected rat lungs was detected by transmission electron microscopy (TEM; Figure 1). These vesicles were distributed throughout the alveolar lumina, surrounding clusters of trophic cells and asci. EVs were also observed between both the host ATI cells and *P. carinii* trophic cells. Additionally, EVs were seen within concave folds of trophic cells, as well as between individual organisms. TEMs of EVs from infected rat lung BALF revealed a heterogenous size population of EVs ranging from 49-266 nm (Figure 2A). The more sensitive nanoparticle tracking analysis showed BALF EVs isolated from uninfected and infected, immunosuppressed rats produced EVs mostly between 100 and 300 nm (Figure 2B-C). These vesicles contained host mammalian proteins CD9 and TSG101, which are commonly used as EV markers for membrane and cytoplasmic proteins, respectively (Figure 2D).

**Figure 1.**
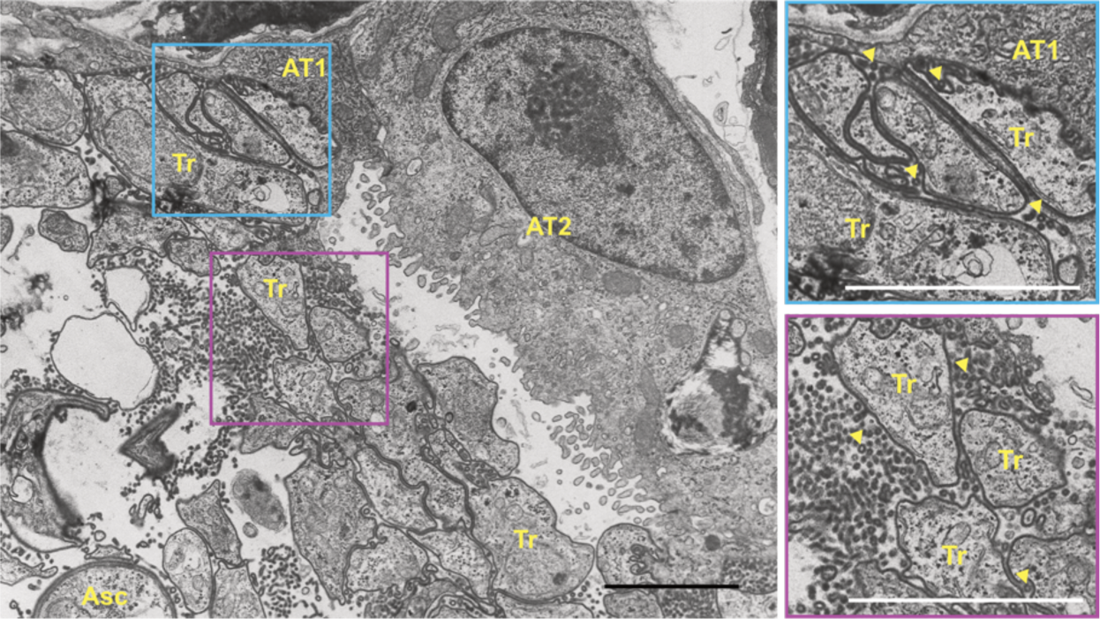
*P. carinii*-infected rat lungs contain abundant EVs. Trophic (Tr) forms were seen within the alveolar space and tightly adhered to the alveolar epithelial type I (AT1) cells. Trophic and asci (Asc) were not adhered to type II cells (AT2). Large population of EVs (arrowhead) were seen surrounding the trophic forms. Colored inset magnified on right. Scale, 10 μm.

**Figure 2.**
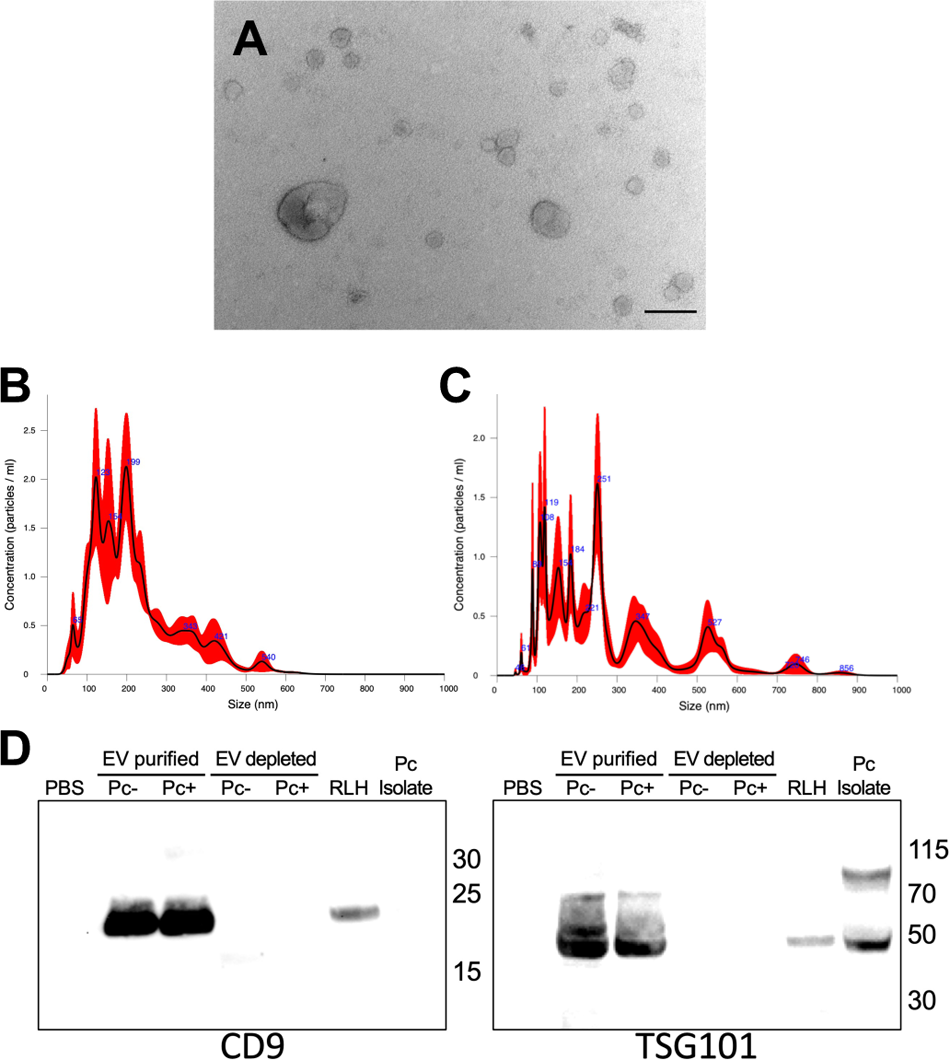
BALF EVs purified from *Pneumocystis carinii*-infected rat lungs. (A) TEM of purified BALF EVs from *P. carinii* infected rats. Scale, 200 nm. (B) Nanoparticle tracking analysis of uninfected rat BALF EVS reveal purified vesicles range from 65-540 nm. (C) *P. carinii*-infected BALF EVs range from 45-856 nm. (D) Western blot of EV marker proteins in BALF EVs from uninfected (Pc-) and *P. carinii*-infected (Pc+) rat BALF. Rat lung homogenate (RLH) from *P.carinii*-infected lungs. CD9 is a tetraspanin membrane protein. TSG101 is a cytoplasmic EV protein.

### P. carinii and P.murina EV proteins

Characterizing the EV proteome derived from *Pneumocystis* spp. presents a challenge, as these fungi cannot be cultured and must be obtained from their host animal. In this study, we sought to identify the fungal EV proteome from infected host BALF by analyzing rat or mouse BALF EVs from immunosuppressed animals using nLC-MS/MS.

*P. carinii* proteins were detected in infected (Pc+) rat BALF EVs (Table 1). These *P. carinii* proteins shared homology with the EV proteomic profiles of other fungal species, such as *A. fumigates*, *C. albicans*, *C. neoformans*, *H. capsulatum*, and *S. cerevisiae* (13, 14, 24–26). Many of the proteins were Heat Shock Proteins, which are commonly used as EV markers for mammalian vesicles and are shown to be contained within the fungal EV proteome (27). This further supports the hypothesis that *P. carinii* secretes EVs.

**Table 1.**
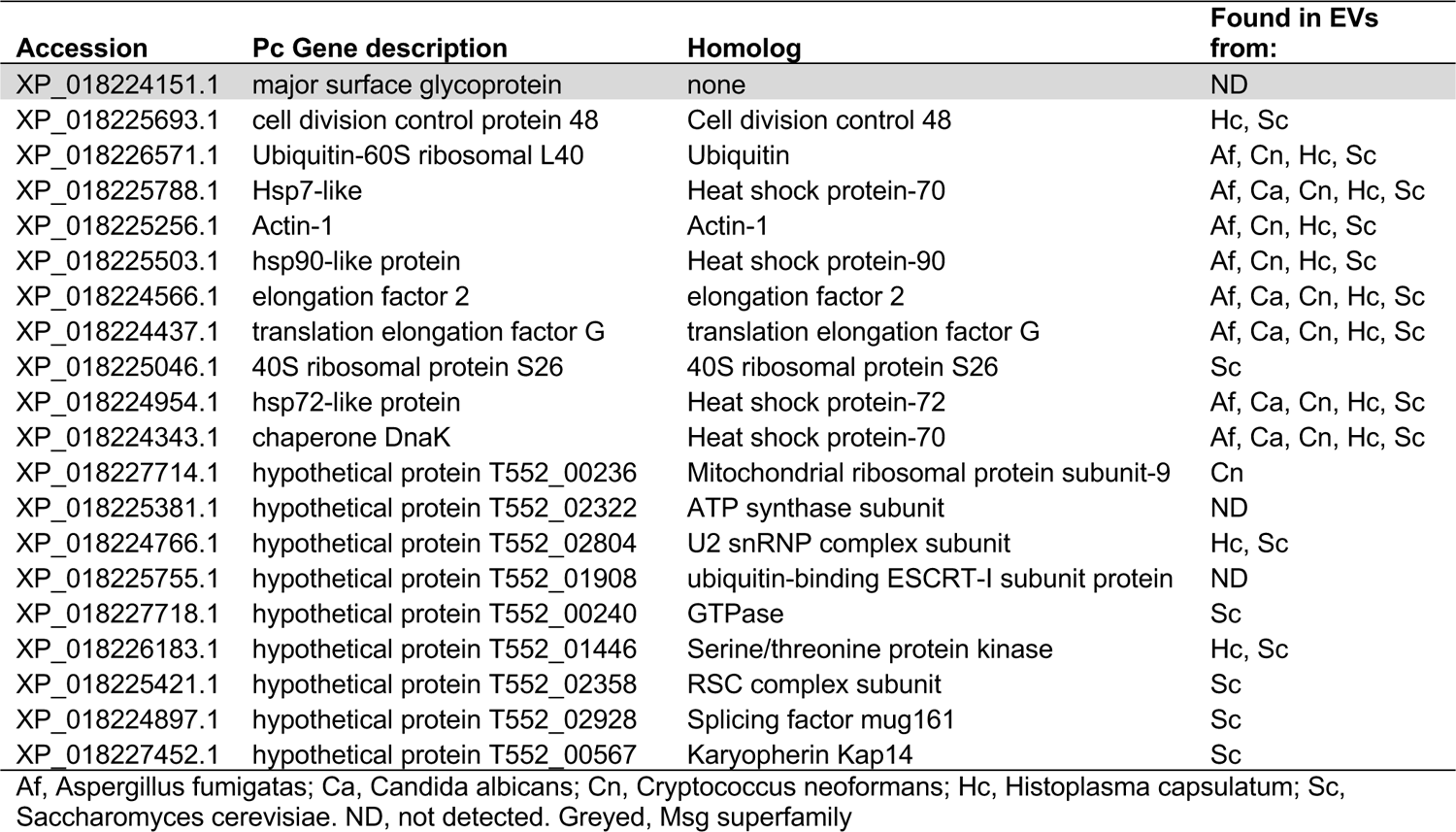
*Pneumocystis carinii* EV proteomic profile from *P. carinii-*infected rat lungs identified by nLC-MS/MS analysis.

Similarly, we detected *P. murina* proteins in infected (Pm+) mouse BALF EVs (Table 2), which also showed homology to the EV proteome of other fungal species. Notably, both *P. carinii* and *P. murina* EVs contained Major surface glycoprotein (Msg). Msg proteins are a superfamily of membranous proteins unique to the *Pneumocystis* genus and are found on the surface of all life cycle stages of these fungi.

**Table 2.** *Pneumocystis murina* EV proteomic profile from *P. murina-*infected mouse lungs identified by nLC-MS/MS analysis.

| Accession | Pm Gene description | Homolog | Found in EVs from: |
| --- | --- | --- | --- |
| XP_007875521.1 | Major surface glycoprotein | none | ND |
| XP_007875520.1 | Major surface glycoprotein | none | ND |
| XP_007875542.1 | Major surface glycoprotein | none | ND |
| XP_007875522.1 | Major surface glycoprotein | none | ND |
| XP_019613321.1 | Major surface glycoprotein | none | ND |
| XP_007874420.1 | Major surface glycoprotein | none | ND |
| XP_007875037.2 | Major surface glycoprotein | none | ND |
| XP_007875038.2 | Major surface glycoprotein | none | ND |
| XP_007873850.1 | Glyceraldehyde-3-phosphate dehydrogenase | Glyceraldehyde-3-phosphate dehydrogenase | Af, Ca, Cn, Hc, Sc |
| XP_007873623.1 | Protein BMH1 | 14-3-3 BMH1 | Af, Cn, Hc, Sc |
| XP_007874173.1 | Actin-1 | Actin-1 | Af, Cn, Hc, Sc |
| XP_007873819.1 | hypothetical protein PNEG_03614 | Metallopeptidase | Cn, Hc, Sc |
| XP_007873020.1 | hypothetical protein PNEG_01105 | Metallopeptidase | Cn, Hc, Sc |
| XP_007875395.1 | hypothetical protein PNEG_03319 | Heat shock protein-90 | Af, Cn, Hc, Sc |
| XP_007871898.1 | hypothetical protein PNEG_00045 | 1,3-beta-glucanosyltransferase | Af, Ca, Cn, Hc, Sc |
| XP_007874319.1 | hypothetical protein PNEG_02319 | None | ND |

Af, *Aspergillus fumigatus*; Ca, *Candida albicans*; Cn, *Cryptococcus neoformans*; Hc, *Histoplasma capsulatum*; Sc, *Saccharomyces cerevisiae*. ND, not detected. Greyed, Msg superfamily

The presence of homologous proteins from *P. carinii* and *P. murina* in BALF EVs from infected animals, along with the identification of Msg membrane proteins, suggests that both species produce and secrete EVs.

### Host proteins from rat and mouse BALF EVs

BALF EVs from immunosuppressed rodents, both uninfected (UI) and infected, were subjected to nLC-MS/MS analysis. The results showed that UI rat BALF EVs contained 408 proteins, while infected Pc+ rat BALF EVs contained 261 proteins. Moreover, UI mouse BALF EVs had 355 proteins, and infected Pm+ mouse BALF EVs had 464 proteins. PANTHER tools (28) were used to assign the functional classifications of these proteins, resulting in the identification of Gene Ontology terms and biological processes (Table 3; Supplementary Table 1).

**Table 3.** Functional enrichment of biological processes within uninfected and infected rat- and mouse-derived BALF EVs. Data represents percent of genes matched against the total number of process hits.

| Biological Process | Rat-derived BALF EVs |  | Mouse-derived BALF EVs |  |
| --- | --- | --- | --- | --- |
|  | Uninfected | Pc-infected | Uninfected | Pm-infected |
| Cellular Process (GO:0009987) | 25.3 % | 19.6 % | 31.1 % | 26.1 % |
| Reproductive Process (GO:0022414) | 0.2 % | 0.4 % | 0.1 % | 0.1 % |
| Localization (GO:0051179) | 10.6 % | 9.1 % | 13.1 % | 11.2 % |
| Interspecies Interaction (GO:0044419) | 1.6 % | 2.2 % | 1.0 % | 2.4 % |
| Reproduction (GO:0000003) | 0.2 % | 0.4 % | 0.1 % | 0.1 % |
| Biological Regulation (GO:0065007) | 13.5 % | 12.2 % | 13.8 % | 12.6 % |
| Response to Stimulus (GO:0050896) | 11.5 % | 13.9 % | 9.7 % | 12.7 % |
| Pigmentation (GO:0043473) | 0.4 % | 0.0 % | 0.0 % | 0.0 % |
| Signaling (GO:0023052) | 5.1 % | 3.8 % | 5.9 % | 5.2 % |
| Developmental Process (GO:0032502) | 2.4 % | 3.1 % | 3.1 % | 2.1 % |
| Rhythmic Process (GO:0048511) | 0.0 % | 0.0 % | 0.1 % | 0.1 % |
| Multicellular Organismal Process (GO:0032501) | 2.8 % | 4.0 % | 3.2 % | 2.5 % |
| Locomotion (GO:0040011) | 0.6 % | 1.1 % | 1.8 % | 1.2 % |
| Biological Adhesion (GO:0022610) | 0.9 % | 1.1 % | 1.3 % | 1.5 % |
| Metabolic Process (GO:0008152) | 17.3 % | 18.3 % | 13.4 % | 15.0 % |
| Growth (GO:0040007) | 1.1 % | 1.3 % | 0.4 % | 0.5 % |
| Immune System Process (GO:0002376) | 6.4 % | 9.5 % | 1.8 % | 6.8 % |

In both animals, the uninfected EV proteome was found to be enriched in Cellular Processes (GO:0009987). While in infected hosts, the BALF EV proteome showed enrichment in Response to Stimulus (GO:0050896) and Immune System Processes (GO:0002376).

### P. carinii actively uptake EVs

For the remaining functional assays, BALF EVs derived from rats were used due to the higher amount of EVs obtained from the lungs compared to mice. EVs from uninfected (UI) and *P. carinii*-infected (Pc+) rats were labeled with the lipophilic membrane dye, PKH26, and incubated with fungal cells (Figure 3). *P. carinii* incubated with PKH26-treated PBS shows no micelle formation of the dye (negative control; Figure 3 first and third row). *P. carinii* treated with PKH26-labeled UI EVs shows bright red punctate staining within fungal clusters (Figure 3 fourth row). No red staining was observed in heat killed (Δ80°C) *P. carinii* (Figure 3 second row). These results indicate EVs uptake by *P. carinii* is an active process, which is lost upon cell death.

**Figure 3.**
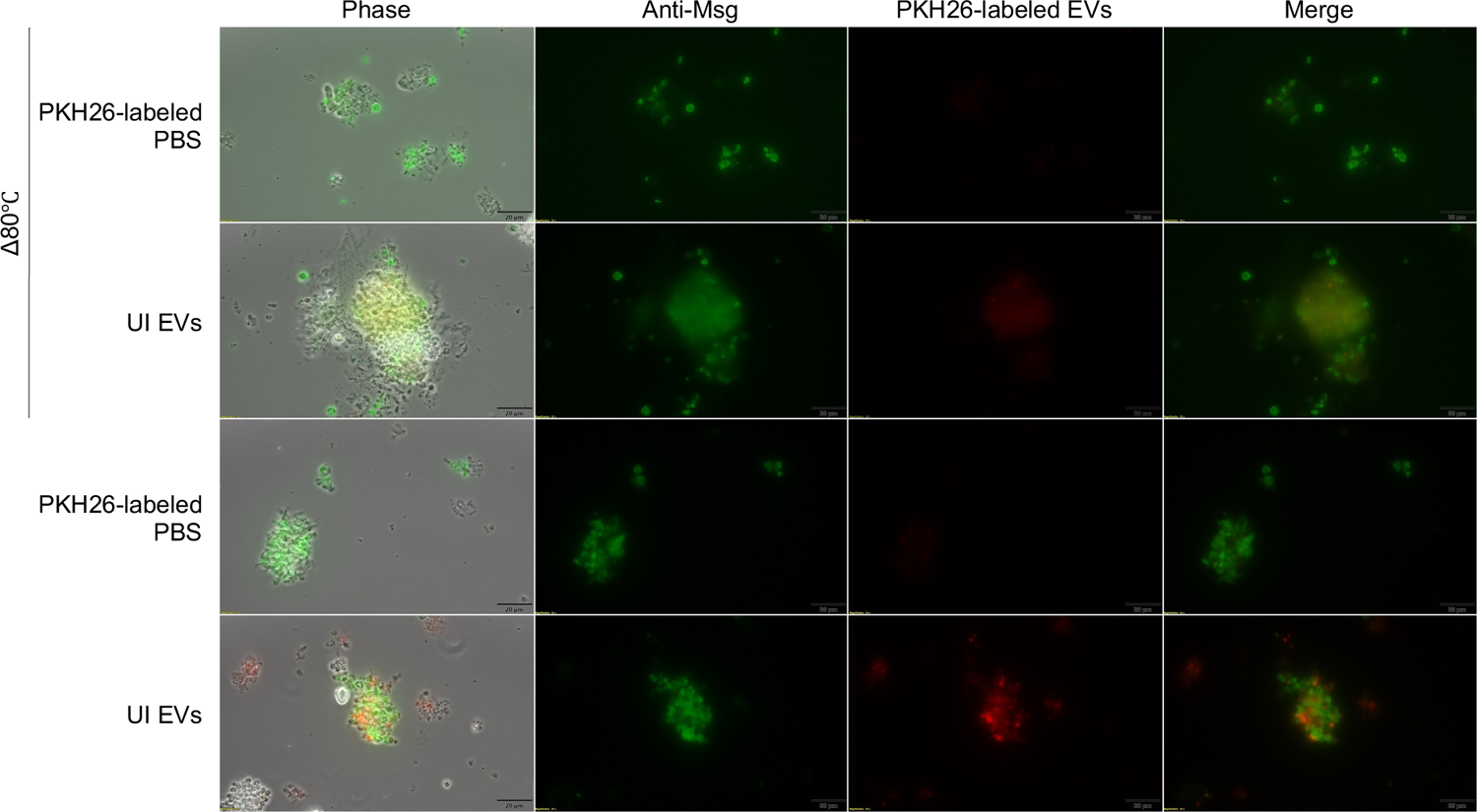
*P. carinii* actively uptake BALF EVs. PKH26-labeled EVs (2 μg) were incubated with either 2.5×106 live or heat killed (Δ80°C) *P. carinii* for 16 hours. Cells were fixed and probed with antibodies directed against Msg. No PKH26 micelle formation was seen in PBS-labeled samples. Δ80°C *P. carinii* treated with PKH26-labeled UI EVs showed no uptake. Live *P. carinii* treated with PKH26-labeled UI EVs shows bright punctate staining within clusters. Scale, 20 μm.

When organisms were treated with Pc+ EVs, *P. carinii* displayed ubiquitous PKH26 staining in both live and dead *P. carinii* cells (Figure 4), indicating that BALF EVs from infected animals are binding to fungal cells irregardless of viability. This is likely due to EV secretion changes during infection and inflammation, which is supported by the Pc+ EV proteome. These EVs contain immune proteins which may bind pathogen-associated molecular patterns on the surface of *P. carinii*.

**Figure 4.**
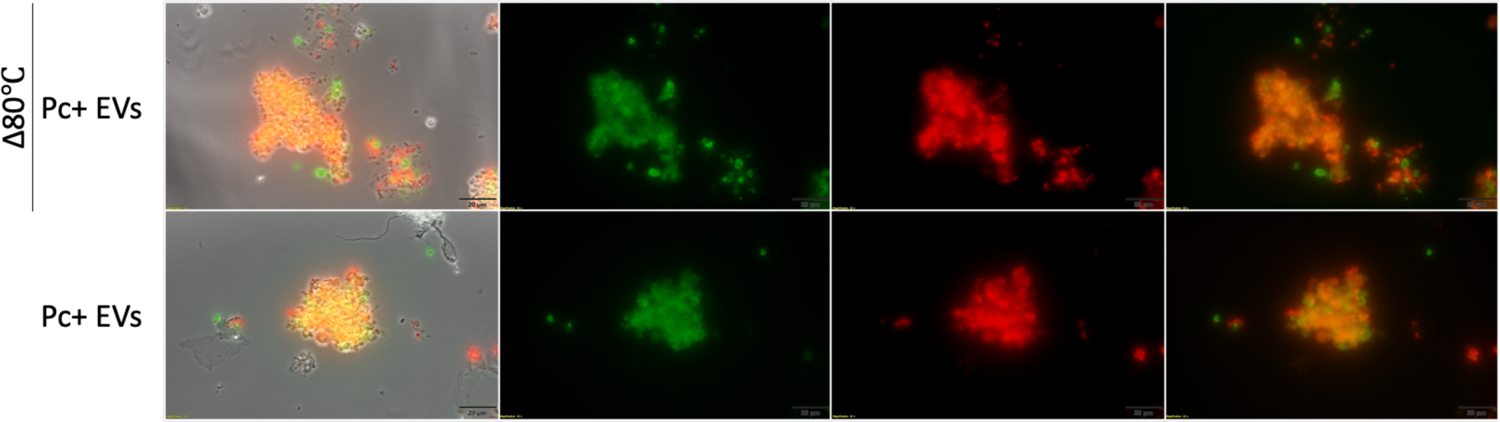
BALF EVs from Pc-infected (Pc+) mice bind ubiquitously to live and dead *P. carinii*. PKH26-labeled Pc+ EVs (2 μg) were incubated with either 2.5×106 live or heat killed (Δ80°C) *P. carinii* for 16 hours. Live and Δ80°C *P. carinii* shows bright staining on the surface of *P. carinii*. Scale, 20 μm.

### Pc+ EVs are detrimental to P. carinii viability

*P. carinii* was treated with EVs from from UI or Pc+ rats and assessed for viability over 7 days (Figure 5A). UI EVs had no effect on viability compared to vehicle and negative control, ampicillin, which is used to test for bacterial growth. However, Pc+ EVs had a detrimental effect on viability, similar to that of the antifungal pentamidine.

**Figure 5.**
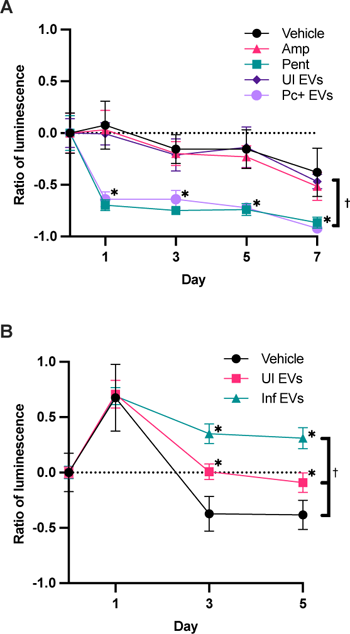
BALF EVs do not improve growth of *P. carinii*. Luminescence data is represented as luminescence compared to the Day 0. (A) *P. carinii* were treated with controls or EVs (2 ug protein equivalent). ATP levels were detected by luminescence using ATPlite. EVs from Pc-infected (Pc+) rats were detrimental to *P. carinii* viability. (B) Rat lung epithelial cells (RLE-6TN) were treated with vehicle or EVs. EVs do not have a detrimental effect on host cell viability. Statistical analyses, ANOVA. *, p<0.05 compared to vehicle at each timepoint; †, p<0.05 compared to vehicle effect.

Since Pc+ EVs were toxic to *P. carinii*, we sought to determine whether these EVs were detrimental to rat lung epithelial cells (RLE-6TN). RLE-6TN displayed no decreased viability when treated with UI or Pc+ EVs. These results indicate while EVs from infected rats were detrimental to *P. carinii* viability *in vitro*, they had no effect on the host epithelial cells.

### EV-stimulated macrophages express pro-inflammatory cytokines

Rat alveolar macrophage cell line, NR8383, were stimulated with EVs from uninfected (UI), infected (Pc+), or zymosan-treated (zymo) rats for 24 hours. Zymosan treatment was used as a positive control to induce a proinflammatory response in the lungs (29). Macrophages treated with any of the EV treatments expressed increased *interlukin (Il)-1β*, *Il-6,* and *tumor necrosing factor alpha (Tnf⍺)* mRNA (Figure 6).

**Figure 6.**
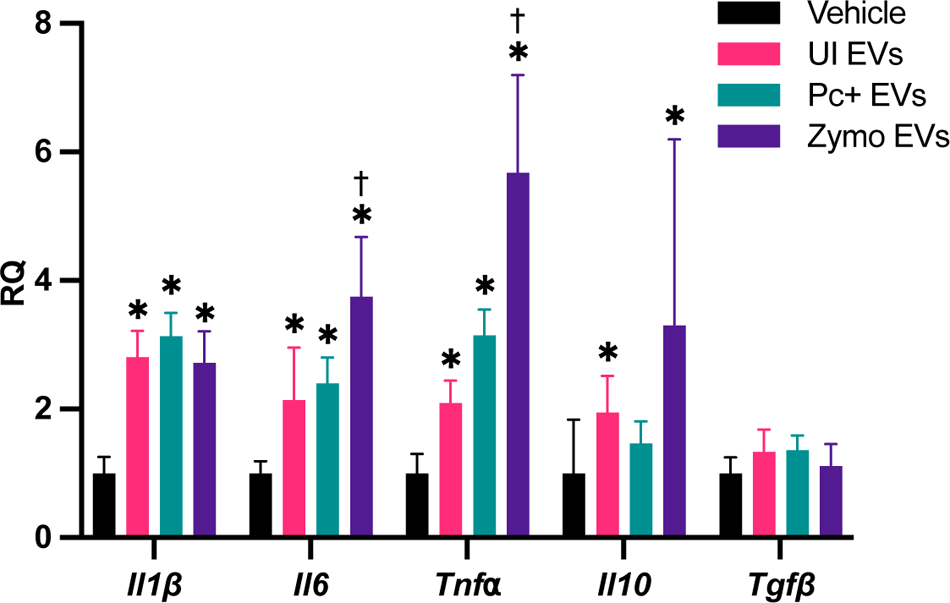
Macrophages stimulated with EVs express pro-inflammatory cytokine mRNA. NR8383 macrophages (4 × 10^4^) were stimulated with vehicle or EVs (2 μg protein equivalent) for 24 hours. EVs from zymosan-treated rats were used as control for a proinflammatory response. Total RNA was extracted from the cells and synthesized to cDNA. RT-qPCR revealed an upregulation of *Il1β, Il6, and Tnf⍺* in EV treated samples, regardless whether they originated from uninfected (UI) or *P. carinii* (Pc+)-infected rats. Statistical analyses, ANOVA. *, p<0.05 compared to vehicle; †, p<0.05 compared to UI EVs.

Macrophages stimulated with zymosan elicited EVs resulted in a more robust expression of *Il-6 and Tnf⍺*, compared to vehicle and UI EVs. However, there were no differences seen between rats treated with UI or Pc+ EVs, indicating the cytokine expression was likely a response to EVs, as a whole, rather than the proteomic differences between UI and Pc+ EVs.

### EVs do not increase macrophage phagocytosis or P. carinii killing

After the NR8383 macrophages were stimulated with vehicle, UI EVs, or Pc+ EVs for 24 hours, *P. carinii* was added to the wells. Macrophages and *P. carinii* were co-cultured for 24 hours to assess macrophage-mediated killing (Figure 7A). Fungal cells were quantified by *PcDhfr* copy number. Macrophages stimulated with EVs, regardless of the origin, did not result in significant killing and quantity of *P. carinii*.

**Figure 7.**
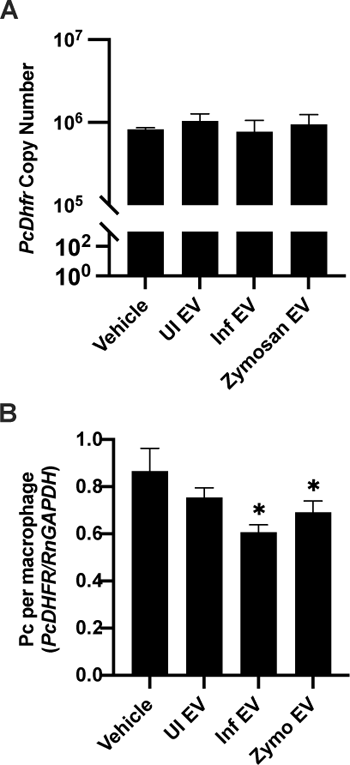
Macrophages stimulated with EVs do not increase *P. carinii* killing or phagocytosis. EVs from zymosan-treated rats were used as control for a proinflammatory response. NR8383 macrophages were stimulated with EVs for 24 hours, then *P. carinii* (2 × 10^6^ nuclei) were added to the wells. (A) To assess macrophages killing *of P. carinii*, wells were collected after 24 hrs coculture. DNA was extracted and fungal organisms were quanitified by qPCR using Pc*Dhfr* copy number compared to a standard curve. (B) Phagocytosis was assessed by collecting the co-culture after 4 hours. Macrophages were washed and differentially centrfigated at 300 x g, then DNA was extracted and used for qPCR. Phagocytosis was quantified by comparing the ratio of *PcDhfr* per *RnGapdh. *, p<0.05 compared to vehicle*.

To assess phagocytosis, vehicle, UI EVs, or Pc+ EV-stimulated macrophages were co-cultured with *P. carinii* for 4 hours (Figure 7B). Macrophages were subjected to differential centrifugation at 300 x g. The ratio of *P. carinii* organisms per macrophage was quantified by qPCR using *PcDhfr* and *RnGapdh.* EV stimulation of macrophages did not significantly increase phagocytosis of *P. carinii* organisms. Perplexingly, phagocytosis was significantly reduced in Pc+ and Zymosan-treated BALF EVs.

## Discussion

Metabolism by *Pneumocystis* spp. is poorly understood, primarily due to the lack of a facilitative culture system (30). Much of our current knowledge of *Pneumocystis* spp. metabolism is based on inferences from genomic studies and coverage of genes from the relevant pathways (5, 31). As with most obligate pathogens, many biosynthetic pathways have been lost, creating a reliance upon the host organism for essential nutrients. This situation poses significant challenges in establishing a suitable *in vitro* culture system and often restricts experimental studies to *in vivo* infection models.

Extracellular vesicles from mammalian tissues and cells known to contain essential components found in *Pneumocystis* spp., such as cholesterol and free amino acids. We posit that *P. carinii and P. murina* take up EVs as a means to supplement their metabolic requirements. To better understand the potential role played by the host EVs, we performed proteomic characterization of EVs from *P. carinii*-infected rat BALF. One challenge with this approach is that the EVs obtained originate from both the rat and the pathogen. As such, we cannot rule out the possibility of host immunity leading to dissociation of fungal cells and presentation of these proteins. However, the identified *P. carinii* EV proteome contains homologues of proteins to those found in the EV proteomes of other fungi including *A. fumigates*, *C. albicans*, *C. neoformans*, *H. capsulatum*, and *S. cerevisiae* (13, 14, 24–26). Additionally, the presence of proteins from the *Pneumocystis*-unique superfamily of membrane proteins (Msg) within BALF EVs suggests that *Pneumocystis* is likely releasing EVs, as vesicles routinely contain membrane proteins from their originating cell. Further research is needed to identify the complete fungal EV proteome from *P. carinii* and *P. murina*. The reasons behind the release of extracellular vesicles by *P. carinii* and *P. murina* remain unclear, but fungal EVs have been shown to serve critical functions in other species. EVs from *C. albicans* and *Pichia fermentans* function in intercellular communication for the proper formation of biofilm (16, 32). Moreover, *Cryotococcus gattii* releases EVs when phagocytosed by host macrophages, leading to long-distance communication and rapid fungal population proliferation (33). These fungal EVs may play important roles in intercellular communication to facilitate growth and survival of *Pneumocystis* within the host lungs.

Host-derived EVs displayed proteins involved in a varity of pathways including cellular and metabolic processes, cellular component organization, and biological regulation. These BALF EVs also include proteins involved in stress response and immune system processes. The specifc host-pathogen interactions of these host vesicles and their cargo with *P. carinii* and *P. murina* are currently unknown.

Neutrophils release EVs in response to *A. fumigatus in vitro* (34). These neutrophil-derived EVs had antifungal properties and inhibited the growth of *A. fumigatus* hyphae. In the case of the phytopathogen *Sclerotinia sclerotiorum*, uptake of host plant EVs caused impaired growth and cell death of fungal cells (35). Additionally, *Arabidopsis* plant cells release EVs containing siRNAs that were taken up by fungal *Botrytis cinerea* cells and induced the silencing of virulence-associated genes (36). Based on these findings, it would be expected that EV host defense mechanisms have a detrimental role in *Pneumocystis* survival within the lungs. This notion is reinforced by the toxic effect observed on *P. carinii* viability *in vitro* when exposed to BALF EVs from *P. carinii*-infected rats.

On the contrary, we suspect that *Pneumocystis* may be also benefitting from the EV contents and utilizing them to supplement their metabolic requirements. Here, we observe host EV uptake by *P. carinii*. This vesicle uptake was lost in heat killed *P. carinii*, demonstrating that EV import is an active process. EV uptake of mammalian-derived EVs may explain how *Pneumocystis* obtains cholesterol from its host. Considering that *Pneumocystis* spp. are deficient in many metabolic pathways, these EVs may also provide amino acids and other necessary components. However, treating *P. carinii* with EVs did not result in sustained growth, or improvement in viability when cultured *in vitro*. This indicates that either EVs are dispensible for *P. carinii* metabolism, or as previously mentioned, *P. carinii* lacks a current sustainable culture system, and the addition of EVs alone is not sufficient to overcome the limitations of this *in vitro* system.

Macrophages participate in host defense response and the clearance of various microorganisms. EVs released by neighboring cells can stimulate these macrophages, leading to their activation and polarization, ultimately promoting inflammation (37). When RAW 264.7 macrophages were stimulated with *A. fumigatus* EVs, they displayed increased killing and phagocytosis of *A. fumagatus* conidia (14). Similarly, *A. flavus* EVs induced M1 polarization of bone marrow-derived macrophages and production of TNF⍺, IL-6, and IL-1β, resulting in increased phagocytosis and killing of *A. flavus* condia *in vitro* (38). In this study, NR8383 macrophages expressed increased mRNA levels of pro-inflammatory cytokines *Il-1β*, *Il-6*, and *Tnf*⍺ when stimulated with EVs, regardless of whether they originated from uninfected or *P. carinii-*infected rat BALF. Additionally, macrophages stimulated with EVs did not exhibit increased killing or phagocytosis of *P. carinii* organisms. These findings are in contrast with macrophages stimulated with fungal EVs. However, our EVs contain a mixture of host, and potentially *P. carinii*, EVs which may explain the contrasting response.

This study reveals the complex interactions between *P. carinii* and host extracellular vesicles (EVs). Host-released EVs have a detrimental impact on *P. carinii* survival within the lungs, while also potentially benefiting the pathogen by supplementing its metabolic needs. Further research is needed to fully understand these interactions and their implications for *P. carinii* survival and metabolism within the host. This area of study shows promise for advancing our understanding of host-pathogen dynamics and exploring potential therapeutic approaches for *Pneumocystis*-related infections. Based on the observed EV uptake by *P. carinii*, it is conceivable that EVs could be packaged with transgenic nucleotide sequences, enzymes, or therapeutic agents for delivery.

## Materials and Methods

### Animals

Sprague Dawley rats were infected with *P. carinii* as previously described (39). In short, Sprague Dawley rats (125-150 g) were housed under barrier conditions with autoclaved food and bedding sterilized in cages equipped with sterile microfilter lids. Rats were immunosuppressed by weekly subcutaneous injections of methylprednisolone [4 mg/kg] and intranasally dosed twice over two weeks with 2×10^6^ organisms of *P. carinii*. Animals continued with 9 weeks of immunosuppression to permit the development a heavy fungal burden.

Corticosteroid-immunosuppressed C3H/HeN mice were infected with *P. murina* through exposure to *P. murina*-infected and immunosuppressed mice. Five week old mice were immunosuppressed by adding dexamethasone [4 µg/L] to acidified drinking water and housed with the seed mice for 2 weeks. A fulminate infection was developed over 6 weeks.

*Pneumocystis* infection was confirmed and quantified by homogenization of the lung tissue and stained with Diff-Quik. The animals were monitored daily, and those showing signs of cachexia were euthanized using an approved method by the AVMA Panel on Euthanasia. These studies were conducted following the guidelines outlined in the 8th edition of the Guide for the Care and Use of Laboratory Animals (National Academies Press, Washington, DC, USA) and in AAALAC-accredited laboratories under the supervision of veterinarians. Additionally, all procedures adhered to the regulations set forth by the Institutional Animal Care and Use Committee at the Veterans Affairs Medical Center, Cincinnati, OH, USA.

### EV purification

Rats and mice were sacrificed after 9 weeks and 6 weeks of infection, respectively. BALF was collected by instillation of cold 0.22 μm-filtered phosphate buffered saline (PBS; 10 mL for rats and 1 mL x 3 for mice) into the bronchiole and alveolar space and gently collected. Cellular debris was removed by centrifugation at 3400 x g for 15 minutes.

EVs were purified using previously described methods (40). Briefly, BALF was filtered using 100kDa Amicon Ultra (Millipore, Darmstadt, Germany). The flowthrough was collected as EV-depleted samples. Size exclusion chromatography (SEC) was performed on filtered BALF samples using qEV10 columns and the Automatic Fraction Collector (Izon Science, Medford, MA). Purified BALF EVs were concentrated by centrifugation at 190,000 x g for 2 hours at 4°C, and the pellet was resuspended in PBS. EV particles were quantified by nanoparticle tracking analysis (NTA) using a NanoSight NS300 (Malvern Panalytical, Malvern, UK). EV protein content was measured using Micro BCA Protein Assay Kit (Thermo Scientific, Rockford, IL).

### Transmission Electron Microscopy

Lung samples were prepared for TEM using previously published methods (41). Briefly, lung samples were fixed in 3% gluteraldehyde/3% acrolein in 0.1 M cacodylate buffer (pH 7.3). Specimens were fixed at room temperature overnight, then post-fixed in 2% OsO_4_ in 0.1 M cacodylate buffer at room temperature for 1 hr. After dehydration in acetone, samples were then embedded in ultralow viscosity plastic. Thin sections were stained with uranyl acetate and lead citrate.

SEC-purified EVs were fixed in 3% glutaraldehyde in 0.1 M cacodylate buffer (pH 7.3) for 30 minutes. Samples were visualized by whole mounting vesicles onto a formvar/carbon-coated 200 grid copper mesh and counterstained with uranyl acetate.

### Western blot analysis

Purified EVs were lysed using radioimmunoprecipitation assay (RIPA) lysis buffer. 2 ug of EV protein content were separated on a 4-12% Bis-Tris gel (Thermo Scientific, Rockford, IL), then blotted onto a PVDF membrane. Nonspecific binding was blocked by Starting Block T20 PBS Blocking Buffer (Thermo Scientific, Rockford, IL) for 1 hour followed by primary antibodies targeting EV membrane protein, CD9 (Abcam, Waltham, MA; 1:10000), or EV cytoplasmic protein, TSG101 (Abcam, Waltham, MA; 1:10000) and incubated for 1 hour. Horseradish peroxidase-conjugated anti-rabbit secondary antibody (Invitrogen, Eugene, OR; 1:50000), was applied to the membrane and incubated for 1 hour. Membranes were washed for 15 minutes 3 times between incubation periods with PBS+0.1% Tween 20. Signal was detected using SuperSignal West Femto Maximum Sensitivity Substrate (Thermo Scientific, Waltham, MA) and imaged using an Invitrogen iBright CL1000.

### Mass Spectrometry

Purified EVs were separated on a 4-12% Bis-Tris gel. The following steps were performed in 25 mM ammonium bicarbonate. Sections were excised, reduced with 25 mM dithiothreitol, alkylated with 55 mM iodoacetamide, and digested overnight with 10 ng/µL trypsin. The peptides were extracted and dried, then resuspended in 0.1% formic acid. Each sample was analyzed by nanoLC-MS/MS (Orbitrap Eclipse, Waltham, MA) and the peptides were matched against a *P. carinii+Rattus norvegicus* or *P.murina+Mus musculus* UniProt database (42; accessed 06/2020) using Proteome discoverer v2.4 and the Sequest HT search algorithm (Thermo Scientific, Waltham, MA). Proteomic analysis was performed on 3 independent EV isolations and data was concatenated for coverage of the proteome. PANTHER tools (28; Release 17.0) were used for functional classifications to identify Gene Ontology terms and biological processes.

### Uptake assays

*P. carinii* EVs were used for the remaining functional assays, as described below, due to the quantity of rat BALF EVs obtained compared to those obtained in mice. *P. carinii* EVs were stained with PKH26 Red Fluorescent Cell Linker Kit (Sigma-Aldrich, St. Louis, MO) for 5 minutes, then quenched with 10% BSA. PKH26-labeled EVs were layered onto 0.971M sucrose and centrifuged at 190,000 x g for 2 hr at 4°C to remove excess PKH26 dye. The resulting pellet was resuspended in RPMI 1640 (Gibco, Waltham, MA).

*P. carinii* (2 × 10^6^) were incubated at 37°C 5% CO_2_ for 2 hours to allow cells to resume metabolic function. Dead cells were produced by heat killing the cells at 80°C for 20 minutes and used as control. PKH26-labeled EVs (2 ug protein equivalent) were added to live or dead *P. carinii*. Cells were incubated at 37°C 5% CO_2_ for 24 hours. The cells were washed in PBS and fixed in 3.7% formaldehyde in PBS for 15 minutes. Cells were attached to slides using CytoSpin 2 (Thermo Shandon, Kalamazoo, MI) at 1000 rpm for 10 minutes. Samples were blocked in 5% goat serum for 1 hour, then incubated with anti-Msg antibodies and anti-rabbit-Alexa Fluor 488 for 1 hour each. Cells were washed between incubations with PBS+0.1% Tween 20 three times for 15 minutes. Samples were imaged using Olympus IX83 inverted microscope.

### Viability Assays

To test the effect of EVs on fungal viability, *P. carinii* (5 × 10^7^ nuclei) were inoculated into 48-well plates (Costar 3548, Corning, New York). Vehicle control or EV samples (2 μg protein equivalent) were added to the wells. Plates were incubated at 5% CO_2_, 37°C. At 1, 3, 5, and 7 days, 100 μL samples were transferred to opaque white plates (USA Scientific, Ocala, FL) and assessed for ATP content using ATPlite (Perkin-Elmer, Waltham, MA).

Rat alveolar cell line, RLE-6TN (ATCC# CRL-2300; 4 × 10^4^ cells/well), was inoculated into opaque white plates (USA Scientific, Ocala, FL) and treated with 2 μg EVs. At 1, 3, and 5 days, samples were assessed for ATP content using the methods described above.

### Macrophage stimulation

NR8383 (ATCC# CRL-2192; 4 × 10^4^ cells/well), a rat macrophage cell line, was inoculated into 48-well plates. Vehicle and EV samples were added to the wells. Plates were incubated at 5% CO_2_, 37°C for 24 hrs.

To assess the cytokine response of EVs on macrophages, RNA was extracted from the cells using Trizol. cDNA was generated from RNA using SuperScript IV VILO Master Mix with ezDNase (Invitrogen, Waltham, MA). Expression was measured using PowerUP SYBR Green Master Mix (Applied Biosystems, Waltham, MA). Relative quanitification of *interluken (Il)-1β*, *Il-6, tumor necrosing factor alpha (Tnfa), Il-10, and transforming growth factor beta (Tgfβ)* was compared against *Gapdh*, as housekeeping gene using the ddCt method. Primers: *Il-1β (*GCTTCAGGAAGGCAGTGTCA; CTCCACGGGCAAGACATAGG*); Il-6 (*AACAGCGATGATGCACTGTCA; ACGGAACTCCAGAAGACCAGA*); Tnfa (*TTCTCATTCCTGCTCGTGGC; AACTGATGAGAGGGAGCCCA*); Il-10 (*CCATGGCCCAGAAATCAAGGA; TTGGGTGGCTACAGGGGAAA*); Tgfβ (*GACTCTCCACCTGCAAGACC; GGACTGGCGAGCCTTAGTTT*); Hprt (*CGACCGGTTCTGTCATGTCG; AAACACCTTTTCCAAATCTTCAGCA).

### Macrophage killing and phagocytosis assays

Macrophages were incubated with EVs as described above. After the 24 hr stimulation period, *P. carinii* (2 × 10^6^ nuclei) were added to the wells. To assess the killing of *P. carinii* by macrophages, wells were collected after 24 hrs. DNA was extracted using DNeasy Blood & Tissue Kit (Qiagen). Fungal organisms were quanitified by *P. carinii dihydrofolate reductase* (Pc*Dhfr*) copy number using TaqMan Fast Advanced Master Mix (Applied Biosciences, Waltham, MA) for qPCR on a ABI 7500 Fast Real-Time PCR system (Applied Biosciences, Waltham, MA). Serial dilutions of known *P. carinii* counts were used to generate the standard curve.

Phagocytosis was assessed by stimulating macrophages with EVs as described above. *P. carinii (2 × 10^6^ nuclei)* were added to the wells. After 4 hours, cells were collected then washed and differentially centrifuged at 300 x g for 15 min three times. DNA was extracted and used for qPCR as described above. Fungal organisms were quantified with *PcDhfr*, while macrophages cells were quantified with Rattus norvigus *Gapdh (RnGapdh)*. Primers: *PcDhfr* (GGCCGATCAAACTCTCTTCC; TCCAGAGATTCATTTCGAGTGAT; /56-FAM/TTGCAATTT/ZEN/CGGCCCCTTAAAGGTC/3IABkFQ/); *RnGapdh (*GGATGGCCCCTCTGGAAAG; GGGGTAGGAACACGGAAGGC; /56-JOEN/CCACTGGTG/ZEN/CTGCCAAGGCTGTGGGC/3IABkFQ/).

## Acknowledgments

This work was supported by NIH R01HL146266 and VA I01BX004441. MTC is a Senior Research Career Scientist supported by IK6BX005232 Department of Veterans Affairs. Protein mass spectrometry data were collected and processed in the University of Cincinnati Proteomics laboratory using mass spectrometry systems supported by NIH instrumentation grants (S10RR027015 and S10OD026717). We thank Scott Langevin and Damaris Kuhnell for assistance with the NanoSight analyses.

